# Feral pigeon populations: their gene pool and links with local domestic breeds

**DOI:** 10.1101/2020.01.18.911198

**Authors:** Dimitri Giunchi, Nadia Mucci, Daniele Bigi, Chiara Mengoni, N. Emilio Baldaccini

**Author notes:** D. Giunchi and N. Mucci contributed equally to the manuscript.

## Abstract

*Columba livia* is a wild bird whose domestication has led to a large number of pigeon breeds. The occasional loss or straying of domestic birds determined the origin of feral pigeons, which are now widespread all around the world. In this study, we assumed that the main contribution to feral populations is provided by domestic breeds reared in the same areas. We tested this hypothesis by analysing the variability of 12 microsatellite loci in nine Italian feral populations sampled in areas with different intensities of breeding and selecting domestic breeds. We included in the analysis samples belonging to domestic lineages commonly bred in Italy The pattern of geographic differentiation of feral populations turned out to be rather complex and only partially explained by the geographic distance between populations. This pattern can be understood only when the domestic breeds were included in the analysis. In particular, feral populations located in regions with a long-lasting tradition of pigeon breeding showed a high level of admixture with domestic breeds, in particular with Racing Homer and Piacentino. Ferals from Bolzano, Venice and Sassari were characterized by unique genetic components, almost all of which are not shared by other feral populations and by the considered domestic breeds. Our results further emphasize the complex origin of feral populations which can be properly investigated only by considering the pool of domestic pigeons bred in the considered area and their past and present distribution.

## Introduction

Feral pigeons *Columba livia* are one of the most common inhabitants of cities all around the world, being a virtually cosmopolitan taxon (Lever 1987). The Neolithic domestication of the wild rock dove, dating back to about 6,000 years BC (Sossinka 1982) and the subsequent selection of the various pigeon domestic breeds represent the initial steps of their origin. Indeed, feral pigeons originated from domestic pigeons abandoned or escaped from farms and then settled in urban habitat (Johnston and Janiga 1995). This process began in the Old World and it is still ongoing in almost every place where domestic pigeons were introduced or bred (Lever 1987; Johnston 1994). Synanthropic wild rock doves seem to have contributed only marginally to the constitution of feral populations and only within their original range (Ballarini et al. 1989; Johnston and Janiga 1995).

The different ways pigeons established themselves in European and North-American urban habitats have been reviewed from a historical point of view by (Johnston and Janiga 1995; Haag-Wackernagel 1998; Baldaccini and Giunchi 2006). Given that nearly any domestic breed had (and still has) the potential to contribute to the feral gene pool, at least two main contributions to feral populations have been identified. Dovecotes had been rather widespread in several European countries until the 19^th^ century [e.g. The British Isles (Ritchie 1920; Gompertz 1957); France (Van der Linden 1950); Italy (Giachetti 1894)]. Pigeons breeding in dovecotes could revert to a free life in towns and sometimes they were even forced to leave their dovecotes e.g. during the French Revolution, when large dovecotes owned by aristocrats were destroyed (Van der Linden 1950). Urban dovecotes are now rare in Europe so they do not represent an important source of individuals for current feral populations. The main contribution to feral populations both in Europe and North America probably was and is still represented by homing pigeons that failed to return to their home loft (Goodwin 1960; Simms 1979; Stringham et al. 2012). In particular, Simms (1979) suggested that juveniles homing pigeons lost at the beginning of their training period constituted an important component of feral populations at least in the UK.

In Italy, pigeon breeding has been an embedded activity since ancient times especially in Northern and Central regions, where several domestic breeds have been selected (McNeillie 1976; Bigi et al. 2016). It can be assumed that these local breeds have mostly contributed to the Italian feral populations, possibly with birds (both dovecote and wild individuals) imported mainly from Egypt (from the range of *C. l. gaddi* and *C. l. schimperi* subspecies) and Spain, used in pigeon shooting ranges till the first half of the last century (Ghigi in Toschi 1939).

In recent years, domestic pigeons have been the subject of several genetic investigations, mainly aimed at understanding their relationships and their geographic origins (Stringham et al. 2012; Shapiro et al. 2013; Biala et al. 2015; Bigi et al. 2016; Domyan and Shapiro 2017), whereas feral pigeons, on the contrary, received rather less attention. While a number of the above-mentioned studies on domestic breeds actually included one group of feral pigeons in their analyses (e.g. Stringham et al. 2012; Biala et al. 2015), only two studies were specifically focused on feral pigeons, with the aim of clarifying the pattern of genetic differentiation within and between urban areas (Jacob et al. 2015; Tang et al. 2018). To our knowledge, no study has systematically investigated the degree of influence of different domestic breeds on the genetic composition of feral populations.

In this paper, we hypothesized that the gene pool of feral pigeons living in a specific urban context should at least partially reflect the prevalent breeding activities of domestic breeds in the area, given the low dispersal propensity of ferals (Hetmanski 2007; Jacob et al. 2015). In particular, we would like to test the hypothesis put forward by (Stringham et al. 2012), that almost all feral populations should show strong affinities with racing breeds.

We compared the genetic pattern of nine Italian feral populations distributed in areas with different domestic pigeon breeding traditions with the aim of: 1) characterizing their genetic composition and structure; 2) testing their affinities with a number of domestic pigeons commonly bred in Italy that could have contributed to their actual genetic pool (Bigi et al. 2016). For this reason, we included in the sampling a number of feral populations belonging to the Pianura Padana (Pavia, Reggio Emilia, Modena), where the breeding of racing pigeons is traditionally widespread and where a significant number of Italian breeds originated (Ghigi 1950; Bigi et al. 2016), and to Central (Pisa, Livorno) and Northeastern Italy (Venezia, Treviso, Bolzano), where racing pigeon breeding is rather less common (Fig. 1) and to Sardinia (Sassari) where racing pigeon breeding is not present. Sardinia possibly still hosts colonies of wild rock doves (Ragionieri et al. 1991; Baldaccini et al. 2000) which might have contributed to the gene pool of feral populations in the area.

**Fig. 1.**
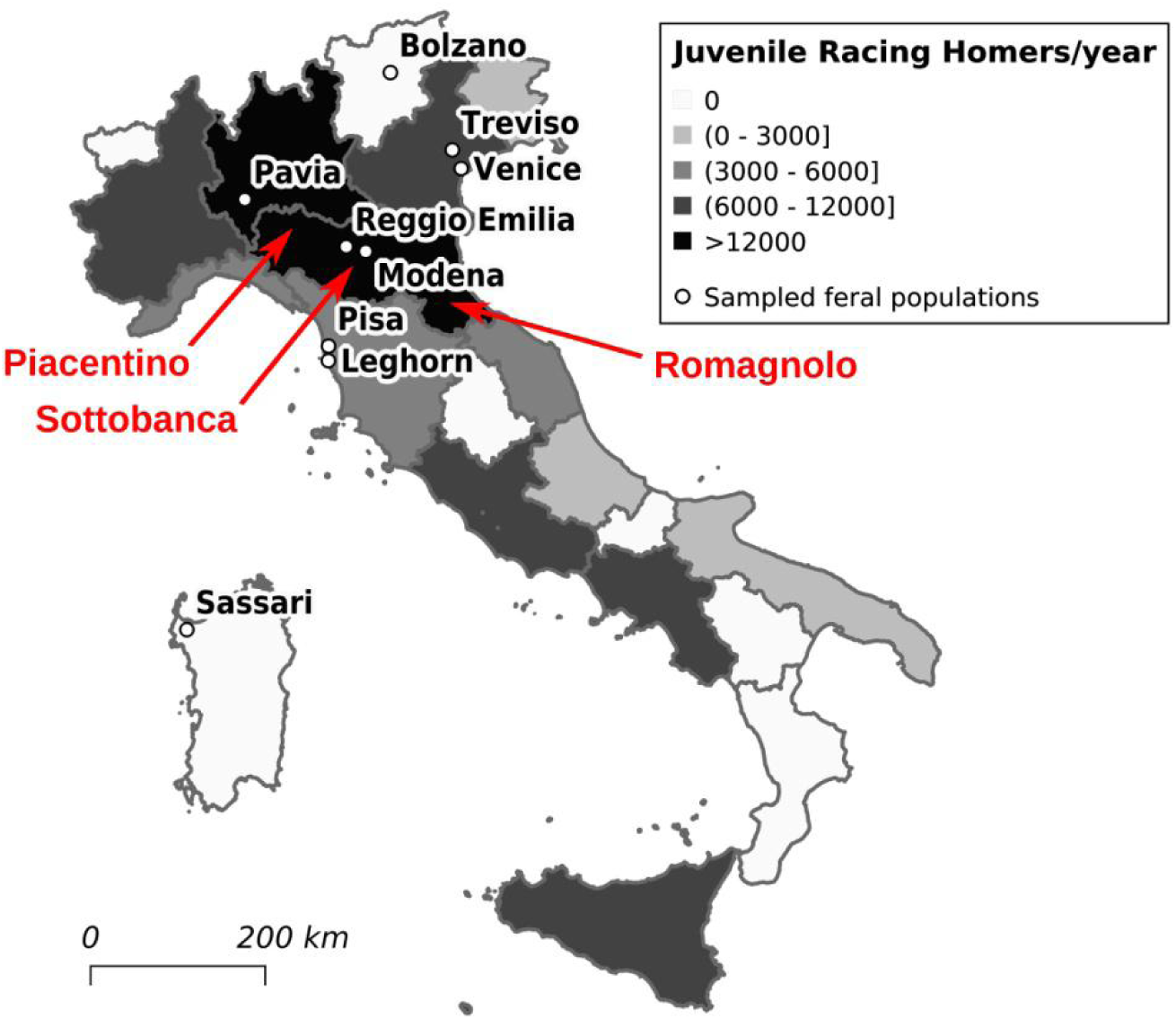
Sampling cities of feral pigeons (white dots), areas of origin of some of the Italian domestic breeds considered in the analysis (arrows) and average number of juvenile Racing Homers reared in the different Italian regions in the period 2013-2018 (data provided by the Federazione Colombofila Italiana). Italian Racing Homer and Runt are not reported in the map because their origin can not be ascribed to a well defined geographical area, even though the selection of the Runt breed probably started in central Italy (Bigi et al. 2016).

## Material and Methods

A total of 194 samples were obtained from nine urban areas (Fig. 1, Table 1) in North and Central Italy. Birds were captured using walk-in traps (Bolzano, Venice, Treviso, Pisa, Reggio-Emilia, Pavia) or were hosted in wildlife rehabilitation centres (Modena, Livorno, Sassari). At least five contour feathers (belonging to the breast and the back) were collected from each individual and stored as soon as possible (usually during the same day of sampling) in ethanol 95% at −20 °C. All the procedures were performed with the permission of the local authorities and complied with the Italian law on animal welfare.

**Table 1.**
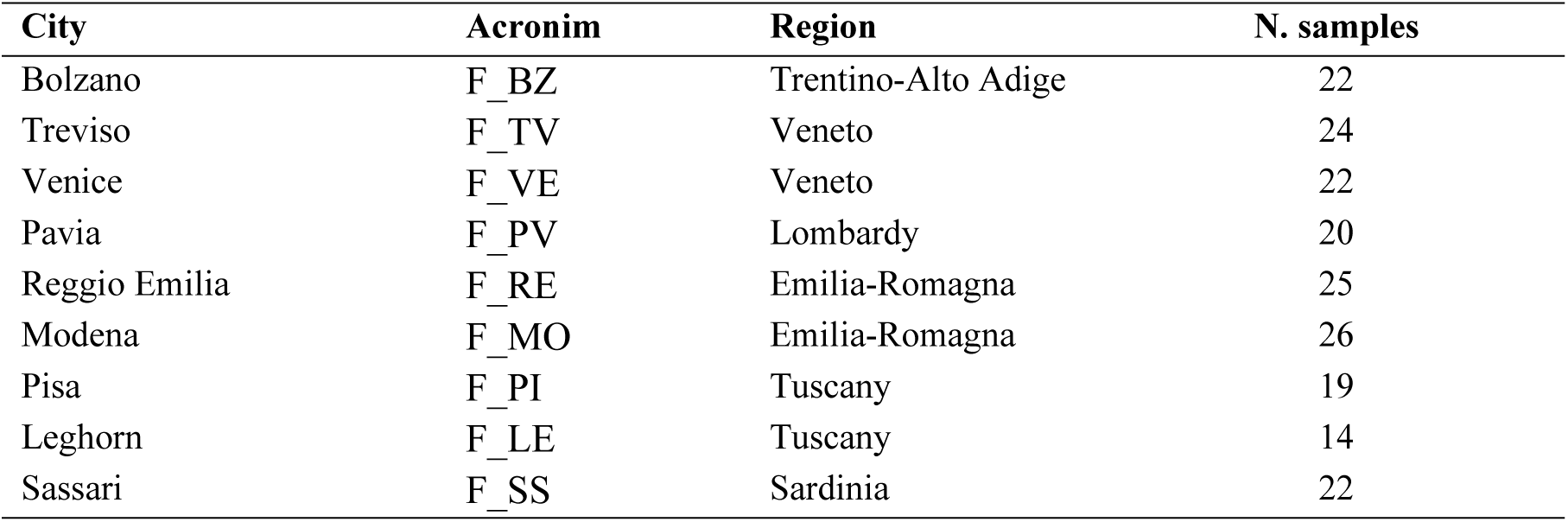
Geographic origin of feral samples.

### DNA extraction and microsatellite genotyping

DNA was extracted using the ZR Genomic DNA II kit (Zymo Research) and amplified at 12 loci microsatellite (*Cliµd01, Cliµt13, Cliµd17, CliµT17, Cliµd16, Cliµd19* - (Traxler et al. 2000); *CliuD11, CliuT47, CliuT24, UUCli10, UUCli13, UUCli14n, UUCli12, UUCli08* - (Stringham et al. 2012). Two independent replicates were performed in 8 μl of volume containing 0.5 U of Hot Start Taq polymerase (Qiagen), 0.18 μM of each primer and 0.04 mg of Bovine Serum Albumin Fraction V (Roche), with the following thermal protocol: (94 °C x 5’), 10 cycles at (94 °C x 40’’) (55 °C-0.5 °C x 40’’) (72 °C x 60’’), 35 cycles at (94 °C x 40’’) (50 °C x 40’’) (72 °C x 60’’), and a final extension at 72 °C for 10’.

### Data analysis

The number of alleles (*Na*), the number of effective (*Ne*) and private alleles, and the expected (*He*) and observed (*Ho*) heterozygosity were obtained using GenAlEx 6.503 (Peakall and Smouse 2006, 2012; Smouse et al. 2015). Allelic richness (*Ar*) was computed in Fstat to minimize the effect of a different sample size (Goudet 2001). Departure from Hardy-Weinberg equilibrium was estimated with the exact test in Genepop on the web (Raymond and Rousset 1995) (http://genepop.curtin.edu.au/) using 100 batches and 1000 iterations per batch.

Pairwise *Fst* computation and AMOVA test (Excoffier et al. 1992) were performed in Genetix 4.05.02 (Belkhir et al. 2002) and GenAlEx 6.503, respectively, to evaluate the significance of genetic differentiation between groups. The matrix of P-values corresponding to each pairwise *Fst* computation was adjusted using the Bonferroni correction (Sokal and Rohlf 1995) for multiple comparisons (nominal level for multiple tests = 0.05).

Genetic divergence among geographical groups was estimated using distance based on the Stepwise Mutation Model (SSM) and the Infinite Allele Model (IAM). Cavalli-Sforza chord distance (*Dc*) (Cavalli-Sforza and Edwards 1967) and the proportion of shared alleles (*Dps*) (Bowcock et al. 1994) were computed in MSA (Dieringer and Schlotterer 2003) and the resulting networks were visualized in SplitTree 4.13.1 (Huson and Bryant 2006). *Da* distance (Nei 1973) and *Fst* distance (Latter 1972) with sample size bias correction, Goldestein distance (*δμ)*^2^ (Goldstein et al. 1995) and Shriver distance (*Dsw*) (Shriver et al. 1995) were computed and visualized in a Neighbour-Joining tree using Poptree on the web (http://poptree.med.kagawa-u.ac.jp/). A total of 10,000 bootstraps were used to reconstruct the tree topology.

Clustering was computed using a Bayesian model in STRUCTURE 2.3.4 (Pritchard et al. 2000; Falush et al. 2003; Hubisz et al. 2009) with no admixture and independent allele frequencies models. The USEPOPINFO selection flag column was considered = 0. A total of five independent runs for a number of subpopulations (K) from 1 to 10, was run with a burn-in period of 30,000 followed by 300,000 MCMC repetitions. STRUCTURE HARVESTER on the web (Dent and von Holdt 2012) was used to process the data and identify the best number of clusters according to both ΔK and Mean Likelihood (Janes et al. 2017). CLUMPP (Jakobsson and Rosenberg 2007) and DISTRUCT (Rosenberg 2004) were used respectively to merge the five independent results for each K and to display the final data. A Discriminant Analysis of Principal Components (DAPC) that is not affected by Hardy-Weinberg disequilibrium was also performed to verify the substructure of feral populations using the Adegenet package (Jombart 2008, Jombart and Ahmed 2011) for R software (R Core Team 2019). A total of 80 PCs and four discriminant functions were retained to draw the plot.

All individuals belonging to the same urban area were characterized by identical coordinates. The hypothesis of Isolation by Distance among populations was tested in GenAlEx by means of the Mantel test calculated on the geographic distance and the *Fst* obtained by GenAlEx computation.

Genetic migration levels between populations was estimated using the function divMigrate of the DiveRsity R pakage (Sundqvist et al. 2016). Nei’s *Gst* and filter_threshold = 0.5 were set to plot the network and the relationships among populations.

### Relationship between feral pigeons and domestic breeds

The considered domestic samples were the same already analysed in (Bigi et al. 2016) and were used to verify the affinity of feral populations with Italian domestic breeds. In details, a total of 250 samples belonging to 10 Italian native breeds (Florentine, Italian Beauty Homer, Italian Owl, Italian Owl Rondone, Piacentino, Romagnol, Runt, Sottobanca, Triganino Schietto and Triganino Gazzo) and one international breed (Racing Homer) commonly bred in Italy, were included in the analysis (Table 2, see Bigi et al. 2016 for further details). *Dc, Dps, Da, Fst*, (*δμ)*^2^ and *Dsw* were computed as for the previous analysis on feral populations to identify the phylogenetic affinities. Cluster analysis was conducted in STRUCTURE using the models applied in the previous section with K ranging from 1 to 20. DAPC was calculated using the same number of PCs and discriminant functions used in the previous section. Cluster analysis and DAPC were run as reported in the previous paragraph.

**Table 2.** Domestic breeds considered in the study. The breeds set off in bold (IO1, IO2, FL, TM1 and TM2) were excluded from the analyses after preliminary investigations (see Material and Methods)

| Domestic Breed | Acronim | No. samples |
| --- | --- | --- |
| **Italian Owl** | **IO1** | **29** |
| **Italian Owl Rondone** | **IO2** | **20** |
| **Florentine** | **FL** | **20** |
| Piacentino | PC | 25 |
| Romagnol | RM | 20 |
| Runt | RN | 19 |
| Sottobanca | SB | 26 |
| **Triganino Schietto** | **TM1** | **26** |
| **Triganino Gazzo** | **TM2** | **20** |
| Racing Homer | RH | 29 |
| Italian Beauty Homer | IH | 16 |

After the preliminary results, five breeds (Florentine, Italian Owl, Italian Owl Rondone, Triganino Schietto and Triganino Gazzo) that did not show any genetic relationship with feral pigeon populations (ESM 1) were excluded from the analysis. The Florentine disappeared in Italy at the beginning of the last century (Giachetti 1894) and it has been reintroduced only recently, while the outdoor breeding of Italian Owls and of Triganino Schietto and Gazzo was interrupted from the first half of 20^th^ century (Vaccari and Zambon 2014). Therefore only a scarce contribution of these five breeds to the present-day Italian feral populations can be expected.

## Results

### Genetic variability of feral populations

The descriptive statistics for the genotypes are listed in Table 3. The allelic number ranged from 6.2 ± 0.5 SE (F_VE) to 8.5 ± 0.9 (F_RE) with an average value of 7.6 ± 0.8. The average number of effective alleles decreased to 4.4 ±0.5. Mean values of expected (*He*) and observed (*Ho*) heterozygosity were 0.718 ± 0.062 and 0.726 ± 0.050, respectively. Allelic richness was computed on a minimum value of 14 samples and ranged from 5.6 (F_VE) to 7.6 (F_LE) with an average value of 6.9. A total of eight different private alleles were detected in Sardinian ferals even if with a low frequency (< 10%).

**Table 3.** Variability indexes. Abbreviations: number of alleles (*Na*), number of effective alleles (*Ne*), allelic richness (*Ar*), observed (*Ho*) and expected (*He*) heterozygosity. Except for *Ar*, values are averages ± SE

| Population | *Na* | *Ne* | *Ar* | *Ho* | *He* |
| --- | --- | --- | --- | --- | --- |
| F_BZ | 6.9 ± 0.7 | 3.9 ± 0.3 | 6.2 | 0.762 ± 0.037 | 0.727 ± 0.027 |
| F_TV | 7.8 ± 0.9 | 4.6 ± 0.6 | 6.9 | 0.705 ± 0.040 | 0.735 ± 0.036 |
| F_VE | 6.2 ± 0.5 | 3.4 ± 0.4 | 5.6 | 0.639 ± 0.065 | 0.651 ± 0.052 |
| F_PV | 7.9 ± 0.7 | 4.6 ± 0.5 | 7.2 | 0.710 ± 0.065 | 0.743 ± 0.038 |
| F_RE | 8.5 ± 0.9 | 4.5 ± 0.5 | 7.1 | 0.739 ± 0.048 | 0.736 ± 0.036 |
| F_MO | 8.3 ± 1.1 | 4.5 ± 0.6 | 7.0 | 0.714 ± 0.056 | 0.724 ± 0.046 |
| F_PI | 8.1 ± 0.8 | 5.0 ± 0.6 | 7.5 | 0.802 ± 0.124 | 0.758 ± 0.036 |
| F_LE | 7.6 ± 0.7 | 4.5 ± 0.6 | 7.6 | 0.708 ± 0.050 | 0.725 ± 0.044 |
| F_SS | 7.3 ± 0.8 | 4.6 ± 0.4 | 6.7 | 0.686 ± 0.069 | 0.733 ± 0.050 |
| Global Mean ± SE | 7.6 ± 0.8 | 4.4 ± 0.5 | 6.9 ± 0.6 | 0.718 ± 0.062 | 0.726 ± 0.050 |

No departure from Hardy Weinberg equilibrium was detected. *Fst* was not significant among feral populations in Central Italy (ESM 2). The highest values were retrieved between Venice and Bolzano (0.093) and between Sassari and Venice (0.081). The lowest values were recorded between Pisa and Treviso (0.016) and between Pavia and Treviso (0.019). AMOVA did not result in a structured sampling with 90% of differences found within individuals and only 4% among populations.

Phylogenetic trees built using *Da, Fst*, and *(δμ)*^*2*^ distances did not show any significant structure (Fig. 2). Ferals from F_LE, F_PI, F_PV, F_MO and F_RE clustered together in all the trees, but the associated bootstrap values were so low as to make the topology not significant. Acceptable bootstrap values were obtained only for the group constituted by F_VE and F_TV. Additional results from other distance computations [Cavalli-Sforza chord distance (*Dc*), proportion of shared alleles (*Dps*) and Shriver distance (*Dsw*)] were not shown as they did not produce a different structure.

**Fig. 2.**
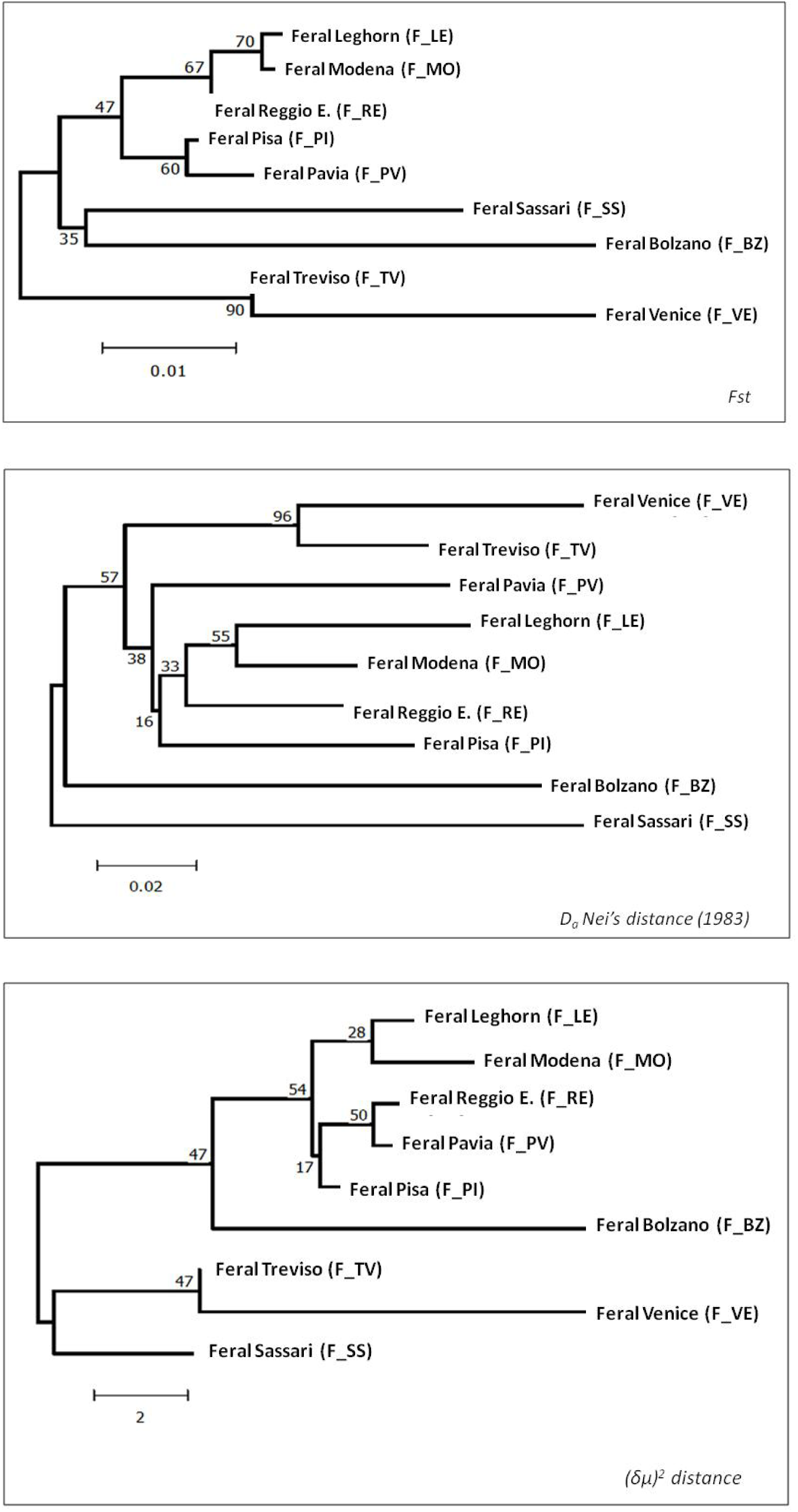
Phylogenetic trees drawn using three different distance-based models: *Da, Fst*, and *(δμ)*^*2*^ distance. No significant topology was supported in any of the three phylogenetic trees. The unique significant clustering was found between F_TV and F_VE.

Mantel test evidenced a positive correlation between the genetic and geographic distances (*r* = 0.50; *P* = 0.03, ESM 3a); this correlation remained almost unchanged (*r* = 0.46; *P* = 0.01) when the Sardinian sample was removed from the analysis (ESM 3b).

No defined genetic structure was identified by both multiv47ariate and Bayesian analyses (Fig. 3a-c). DAPC analysis showed an admixed pattern in Central Italy, whereas samples from Sassari (F_SS), Bolzano (F_BZ), and Venice (F_VE) clustered separately from the others. The population from Treviso (F_TV) plotted in an intermediate position between the remaining populations and F_VE.

**Fig. 3.**
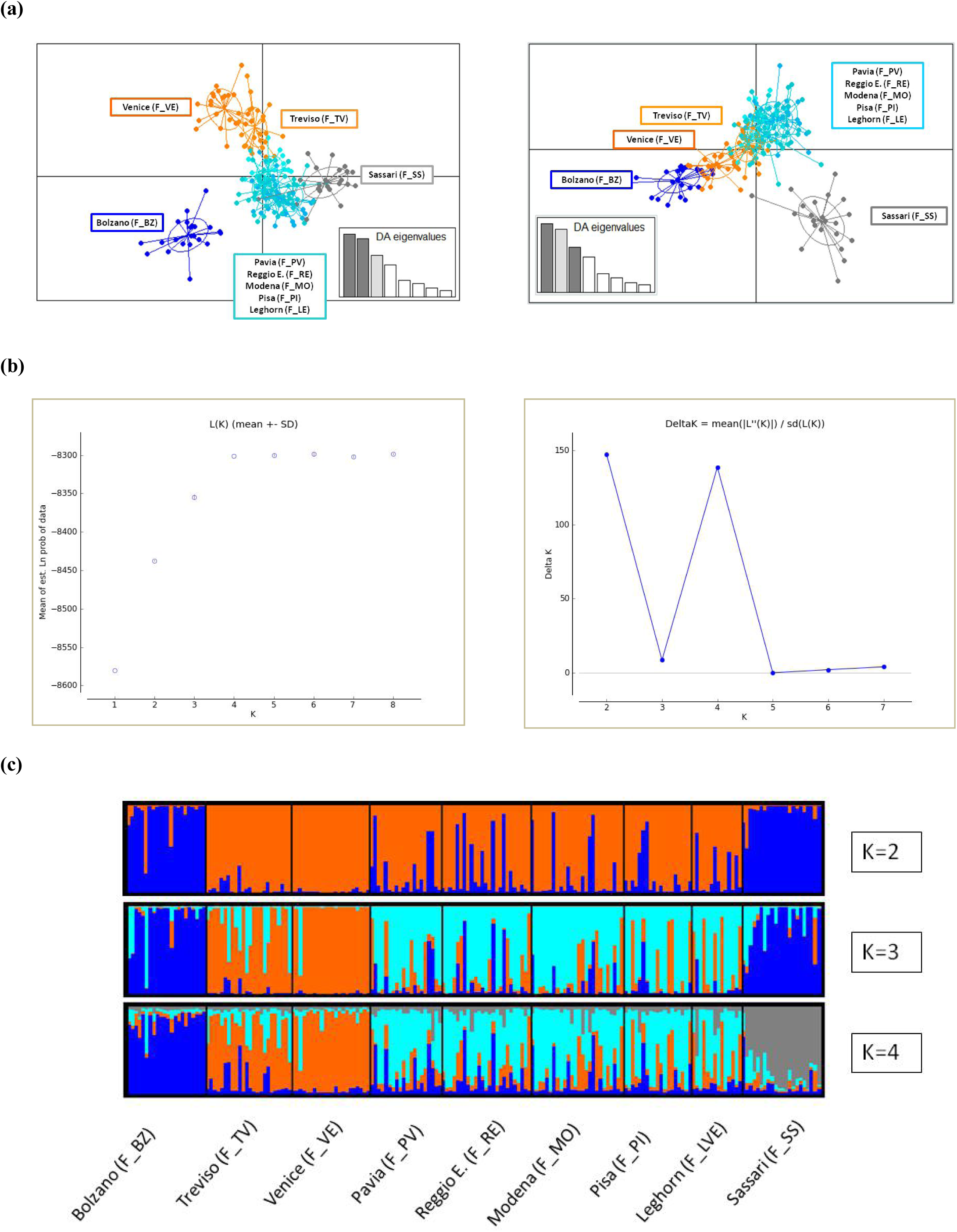
Results of the Discriminant Analysis of Principal Components (DAPC) and of Bayesian computations on feral pigeon samples. **(a)** DAPC evidenced a clear differentiation of F_BZ, F_SS, and F_VE, individuals from Central Italy plotted together, while the position of F_TV was intermediate between F_VE and F_PV. **(b)** The ΔK and mean likelihood computations suggested that 4 clusters represented the best genetic subdivision of the sampling. **(c)** Bayesian analysis assigned F_BZ, F_VE, and F_SS to unique and distinctive populations. At K = 4, the different colored bars in F_TV, F_PV, F_MO, F_RE, F_LE, and F_PI describe admixed genetic compositions within these populations, thus evidencing a relevant genetic flow among them. The barplot at K = 2 and K=3 allows tracking the genetic components and the main subdivisions among groups.

Both Mean Likelihood and ΔK obtained from STRUCTURE HARVESTER identified the best grouping at K = 4. At K = 2 Bolzano and Sassari populations split from the remaining populations. K = 3 permitted an additional distinction of ferals from Treviso and Venice. At the best value K = 4, ferals from Bolzano (F_BZ), Venice (F_VE), and Sassari (F_SS) were associated to different clusters with high *q* individual membership values, while samples from F_PV, F_RE, F_MO, F_PI, and F_LE remained not differentiated from each other and showed retraces of admixture with F_BZ, F_VE, and F_SS. The genetic composition of F_TV was mainly associated with F_VE but showed traces of admixtures with the former five populations (Fig. 3c).

No genetic components shared between Bolzano (F_BZ), Sassari (F_SS) and the other feral populations was estimated by the divMigrate function (Fig. 6). Gene flow towards Treviso (F_TV) from Venice (F_VE) and Pavia (F_PV) resulted well supported (asymmetric values = 0.8 and =0.52, respectively), as well as that from Pavia (F_PV) to Reggio Emilia (F_RE). Populations from Modena (F_MO), Reggio Emilia (F_RE), Leghorn (F_LE) and Pisa (F_PI) resulted to be quite interconnected. The highest values of gene flow (0.94 and 1.0) were reported between Modena (F_MO) and Reggio Emilia (F_RE).

**Fig. 4.**
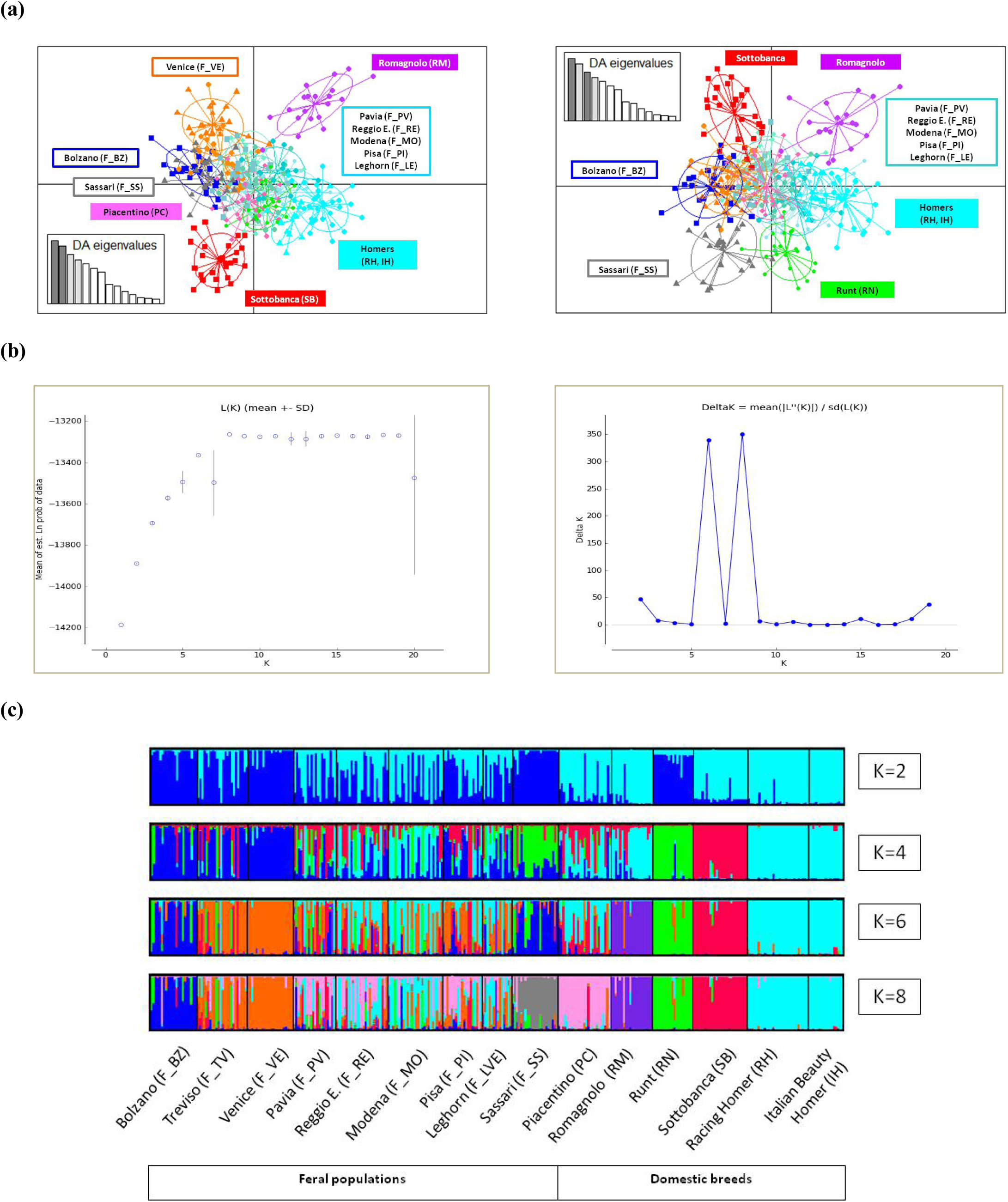
Discriminant Analysis of Principal Components e Bayesian computations in feral and domestic breeds. The computations involved only pigeon lineages that could have contributed to the genetic composition of feral populations. **(a)** DAPC showed that SB, and RM contributed marginally to the sampled feral populations, while IH, RH, PC and in part RN clustered together with ferals from Lombardy, Emilia-Romagna and Tuscany. **(b)** The ΔK and mean likelihood computations suggested that 8 clusters represented the best genetic substructure of the sampling. **(c)** Bayesian analysis evidenced a sharp distinction between feral and domestic samples. At K = 2, the main difference was found between Homers with other domestic and feral pigeons. At K = 4, 6 and 8 the greater subdivisions were internal to both domestic and feral groups. Moreover, light blue, pink colored and light green bars found in ferals at K = 8 suggested a probable origin of some individuals from IH, RH, PC and RN breeds.

**Fig. 5.**
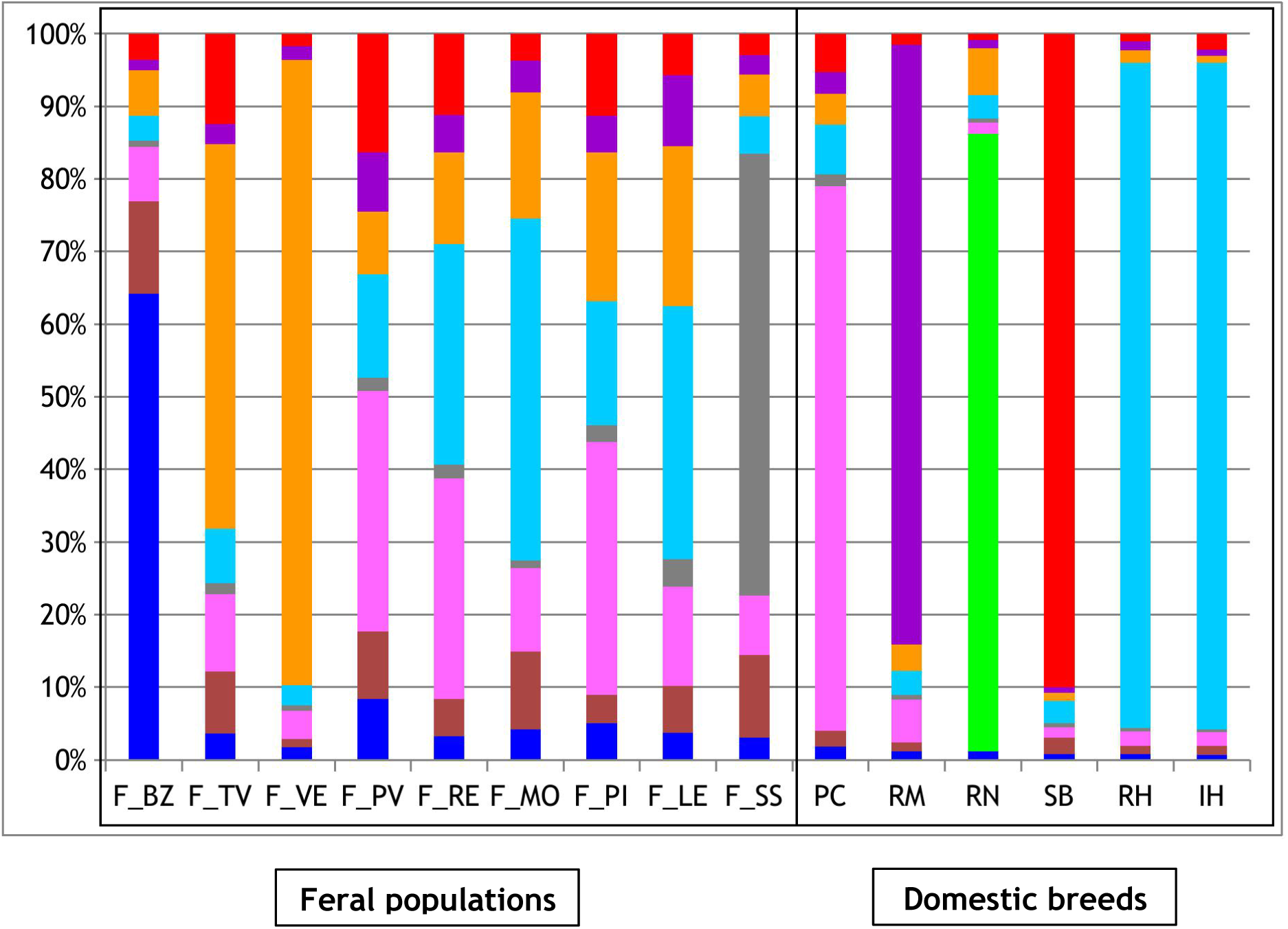
Bar chart showing population membership (Q) values at K=8. Admixed colored bars are representative of an admixed genetic composition and origin. Identical colors indicate a common origin.

**Fig. 6.**
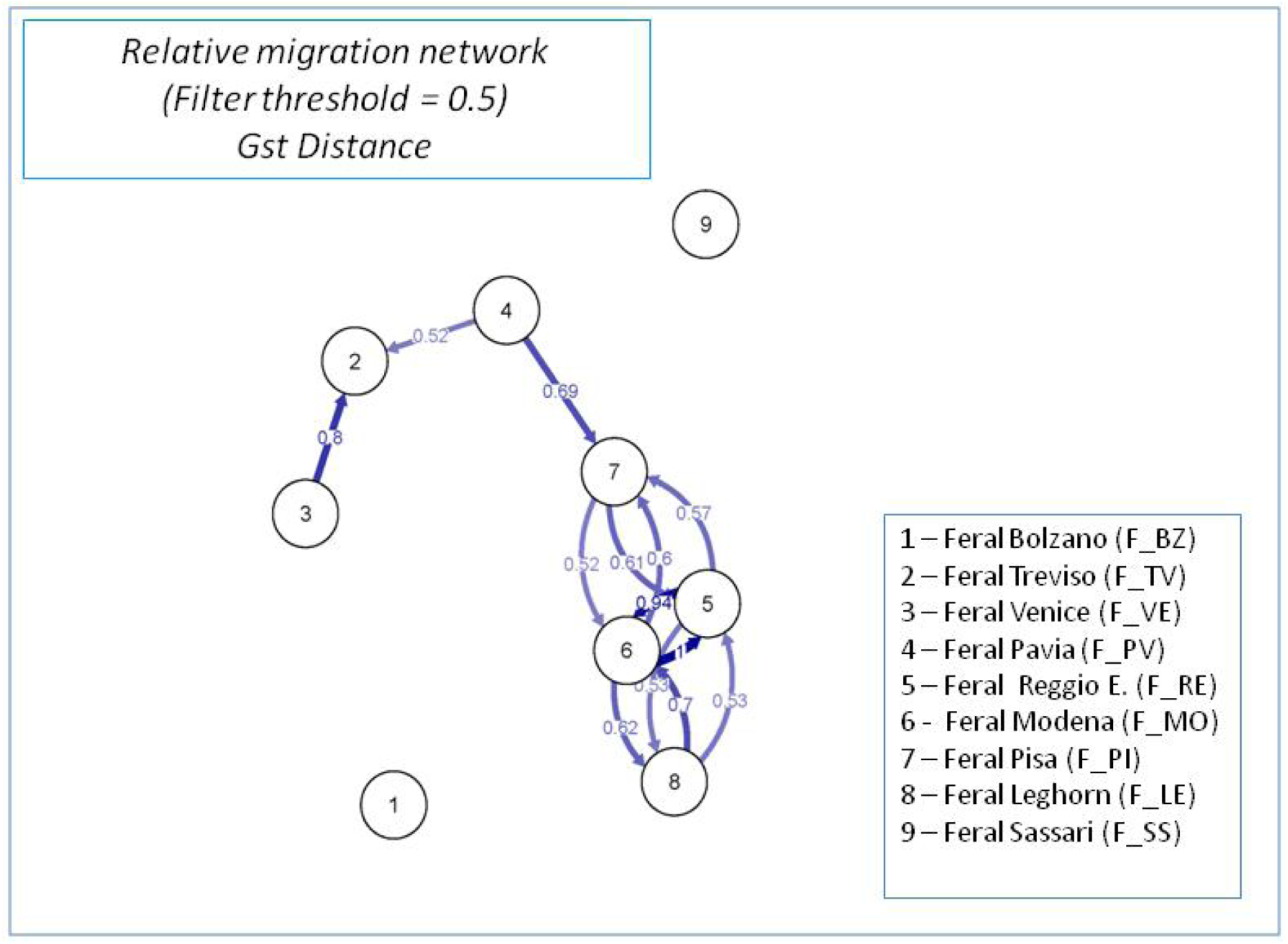
Directional relative migration estimated by divMigrate in DivRsity. The circles represent the feral populations from this study while arrows and numbers identify the direction and the value of migrations. Only significant asymmetric links with values higher than 0.5 were plotted.

### Relationship between feral pigeons and domestic breeds

Phylogenetic trees built using *Fst*, D*a* and *(δμ)*^*2*^ distances did no show any significant structure (ESM 4) and feral groups and the domestic lineages considered in the analysis were differently associated depending on the distance computation considered. As in the previous section, Cavalli-Sforza chord distance (*Dc*), the proportion of shared alleles (*Dps*) and Shriver distance (*Dsw*) were not informative and were not shown.

The DAPC of the nine feral populations and the six Italian breeds did not reveal a sharp separation between the two groups (Fig. 4a). With the exception of domestic SB and RM, all the other breeds overlap with feral populations in the plot. F_VE and F_SS did not show any tracks of an origin from the considered domestic lineages because in the plots they did not overlap with any of them. The Bayesian analysis partially confirmed the results of DAPC and the *q* individual membership values identified a relationship between domestic and local feral pigeons (Fig. 4c and 5). The ΔK computation in STRUCTURE HARVESTER identified 8 main informative clusters (Fig. 4b). The main splits at K = 2, 4 and 6 were also plotted in Figure 4c to describe the main similarities among groups. Interestingly, the first split (K = 2) does not occur between feral and domestic pigeons but between Homers (RH and IH) and the other groups. At K = 4, Runt (RN) and Sottobanca (SB) separated from other domestic lineages while ferals from Sassari (F_SS) separated from peninsular populations. At K = 6, ferals from Sassari, Bolzano (F_BZ), Venice (F_VE) and Treviso (F_TV) separated from the other populations while Romagnolo (RM) splits from Homers. The evaluation of the estimated membership coefficient for each population *(q*) at K = 8 identified the predominance of the domestic component characterizing the Homers (light blue bar in Fig. 5) in feral populations from Lombardy, Emilia-Romagna and Tuscany, particularly in F_RE, F_MO, and F_LE. Rather high percentages of domestic Piacentino (PC) and lower values of domestic Sottobanca (SB) and Runt (RN) were found in all populations, including F_BZ and F_SS, although the latter populations were characterized by over 60% of a unique private component. Ferals from Venice (F_VE) showed a high percentage of a unique genetic component, that was also detected in the other feral populations but that was almost irrelevant in domestic lineages. As evidenced in Figure 5, the population of Venice did not show relevant traces of admixture with the considered domestic populations.

## Discussion

This paper represents one of the few studies dealing with the genetic structure of feral pigeon populations. Up to now, only one study was specifically focused on clarifying the pattern of genetic differentiation of these birds between urban areas (Jacob et al. 2015). Furthermore, while it is largely accepted that feral pigeons originated from domestic breeds (see for instance, Johnston and Janiga 1995), this study is the first one that systematically investigates the affinities between feral pigeon populations and the domestic breeds commonly reared in the same area. To our knowledge only Biala et al. (2015) tried to quantify the gene flow between several domestic breeds and feral pigeons. However, while Biala et al. (2015) used only one feral group, composed of birds sampled in different towns, we sampled a significant number of true feral populations in order to investigate the contribution of domestic breeds to their gene pool and how this contribution varies among populations located in different areas.

On the whole, the populations analyzed in our study showed levels of genetic variability lower than those observed by Jacob et al. (2015). The low levels of variability indices which were found in the populations lacking genetic admixture allowed us to argue that these values were influenced by the reduced incidence of gene flow between these populations and other feral populations.

As observed by Jacob et al. (2015), also in our samples the genetic distance between feral populations was correlated with their geographic distance, both considering all samples or only the peninsular ones. This result further confirms that the exchange rate of individuals among cities is relatively rare (Johnston and Janiga 1995; Hetmanski 2007). It should be noted, however, that the observed pattern of geographic differentiation is rather complex and only partially explained by the geographic distance between populations. Indeed, while the present data do not support a well defined genetic structure, it is interesting to observe that Sardinia and North-eastern populations (F_BZ, F_VE and F_TV to a lesser extent) tended to cluster separately and showed a null or rather low gene flow with the remaining populations. On the contrary, populations belonging to Tuscany, Emilia-Romagna and Lombardy (F_PI, F_LE, F_MO, F_RE and F_PV) showed a high level of admixtures almost independent of their geographic distance. The genetic distinction of populations like F_BZ, F_SS and F_VE, can be attributed to their relative geographic isolation. Actually, Bolzano is located in an Alpine valley surrounded by habitats mostly unsuitable for pigeons, while Sardinia is an island and it is known that pigeons do not like to fly over large water bodies (Wagner et al. 1972). The effect of the surrounding water should probably be taken into account also for Venice: for instance, as reported by (Soldatini et al. 2006), the number of Venetian pigeons involved in foraging flights outside the city is very low considering the size of the population, which possibly confirm the low propensity of these pigeons to fly over the lagoon and thus also to disperse inland. The relatively high gene flow from F_VE to F_TV estimated by divMigrate can be interpreted both considering the relatively short distance between the two cities, but also the likely common origin of the two populations.

The level of admixture of the remaining populations (F_PI, F_LE, F_MO, F_RE and F_PV) and the high level of estimated gene flow among them as well as between F_PV and F_TV are quite difficult to explain considering the above mentioned low rate of dispersal among cities (Johnston and Janiga 1995; Hetmanski 2007; Jacob et al. 2015). and the inability of feral pigeons to undergo long flights, as experimentally demonstrated by Chelazzi and Pineschi (1974) and by Edrich and Keeton (1977). However, it is not to be excluded that any dispersal event could occur between very close cities (e.g. Modena and Reggio Emilia). Actually, the migration events described by DivMigrate are estimated from the allele frequencies retrieved inside the populations and could be interpreted as common genetic components rather than a real gene flow. The inclusion in the analysis of domestic samples confirms this hypothesis. Indeed, in these feral populations it is possible to identify a significant component belonging to the domestic breeds considered in this study. In particular, the Racing Homer and Piacentino components largely dominates the gene pool of these populations. These domestic components are still detectable in F_SS, F_BZ, F_VE and F_TV, but at negligible percentages. This pattern can be explained by considering the long-lasting and still ongoing tradition of keeping and selecting pigeon breeds especially in Emilia-Romagna and Lombardy (Ghigi 1950; McNeillie 1976). In particular, the Racing Homer component is quite evident mainly in populations located in areas where Racing Homer breeding and racing is widespread (i.e. Emilia-Romagna and Lombardy, see Figure 1), while it is almost absent where those activities are missing (i.e. Bolzano and Sardinia). Our data only partially confirm the hypothesis by Stringham et al. (2012, but see also Goodwin 1960; Simms 1979) that Racing Homers constitute the most important component of feral pigeon populations. Indeed, other breeds, such as Piacentino, can be dominant or co-dominant in the feral gene pool, as observed for example in F_PI, F_PV and F_RE.

Our results further emphasize the complex origin of feral populations and suggest a past and probably ongoing flow of domestic pigeons into feral populations in areas surrounded by a high number of pigeon fanciers. This seems to be confirmed by the difference observed between F_VE and F_TV. These populations form a fairly separated cluster, which probably indicates that they share a common origin. However, being located in a lagoon, Venice has no pigeon fanciers nearby and thus the genetic contribution of the domestic breeds considered in this study to its feral population is very low. On the other hand, Treviso is surrounded by farms that probably hosted and still host pigeon dovecotes, which facilitates the urban drift of domestic birds. It should be noted that it is rather impossible to have detailed information regarding the actual distribution of pigeon breeding around a given city and for this reason the above pattern is characterized by some unexplained variability that might be related to the scale of our analysis.

The mechanisms leading to the admixture between domestic and feral pigeons have probably been and still are both pigeon racing and feral pigeon foraging behaviour. As observed by Goodwin (1960) and Simms (1979) pigeon races are sources of numbers of lost Racing Homers that flock together with ferals. Furthermore, the daily foraging flights of ferals towards the surrounding crop fields (Johnston and Janiga 1995; Giunchi et al. 2012) may encourage farm dovecote individuals to join them. As a partial support to this hypothesis, it should be noted that, birds living in cities mostly surrounded by an unsuitable foraging habitat show both less propensity to perform foraging flight [i.e. Bolzano (Baldaccini et al. 2015) and Venice (Soldatini et al. 2006)] and a less relevant component of the studied domestic breeds in their gene pool.

Recent data on the monk parakeet *Myiopsitta monachus* and ring-necked parakeet *Psittacula krameri* suggest that a high degree of admixture is not directly related to invasive success in an urban habitat and does not prevent the possibility of rapid adaptation to the urban environment (Edelaar et al. 2015; Le Gros et al. 2016). In this regard, it would be interesting to a use genomic approach in order to test whether the populations with a higher degree of admixtures, actually show higher frequencies of phenotypic characters belonging to domestic breeds or whether the domestic phenotypes are quickly counter-selected by the urban environment (see Johnston and Janiga 1995; Sol 2008)

As observed above, feral pigeons from Bolzano, Venice/Treviso and Sassari were characterized by unique genetic components, that are mainly not shared by other feral populations. Considering the geographic position of those populations, it can be hypothesized that these unique components belong to domestic breeds not originated and/or reared in Italy (e.g. central-european breeds for Bolzano, eastern breeds for Venice/Treviso). Concerning birds from Sassari, it should be noted that Sardinia still hosts wild populations of rock doves (Ragionieri et al. 1991; Johnston 1992; Johnston and Janiga 1995). Johnston and Janiga (1995) indicated that when wild and feral pigeons live in sympatry it is likely that they interbreed, thus it can be hypothesized that wild pigeons may have contributed to F_SS. In this regard, it is interesting to observe that, contrary to Jacob et al. (2015) and Biala et al. (2015), we found eight private alleles in feral samples, all belonging to the Sardinian population. This result could be explained by considering at least three possible factors: a) the effect of genetic drift, being Sardinian populations relatively isolated (see above); b) the effect of domestic breeds not included in our sample and not affecting other feral populations or c) the gene flow between wild rock doves and feral pigeons as hypothesized by Ragionieri et al. (1991). Our data do not allow to discriminate among these effects so this topic deserves further investigations.

Our data emphasize the critical role of the sampling protocol when studying the relationship between feral pigeons and domestic breeds. Indeed, papers studying this topic often did not consider single populations of feral pigeons, but mixed together pigeons sampled in different cities (see e.g. Biala et al. 2015; Shao et al. 2019). Moreover the domestic breeds included in the analysis were selected without taking into consideration the local tradition of pigeon breeding in the areas where feral pigeons were sampled. This probably explains some of the inconsistencies in the results obtained for instance by Stringham et al. (2012) and Shao et al. (2019), as the former suggested a strong relationship between Racing Homers and feral pigeons, which is not evident in the latter study.

To conclude, our data provide the first detailed analysis of the relationships between feral pigeon populations and domestic breeds, shedding light on the way feral populations originated and are maintained. Our results emphasize the complexity of the feral gene pool whose composition shows high spatial variability possibly depending on both ecological and anthropic factors. In particular, factors such as geographic isolation of feral populations along with the prevalent farming activities and the local diversity of domestic pigeon breeds seem to play a central role in this regard. Further studies are needed in order to investigate the role of wild rock dove on Italian feral gene pool and in particular on Sardinian feral populations.

## Acknowledgments

We thank local regional authorities for providing permits to sample feral pigeons across Italy. We also thanks Andrea Barbon, Pierluigi Berra, Danilo Pisu, Cecilia Soldatini and the staff of the wildlife recovery centres of the following cities for their collaboration in collecting samples: Livorno (Gianluca Bedini, Renato Ceccherelli, Nicola Maggi), Modena (Piero Milani), Reggio-Emilia (Sergio Mazzali), Sassari (Marco Muzzeddu). Thanks are due also to the Federazione Colombofila Italiana and to the Federazione Italiana Allevatori Colombi (FIAC) for samples of domestic breeds and for data regarding the rearing activity of Racing Homers in Italy. Wendy Doherty edited the language of the manuscript.

