## Supplementary Materials for "Feral pigeon populations: their gene pool and links with local domestic breeds"

**(a)**

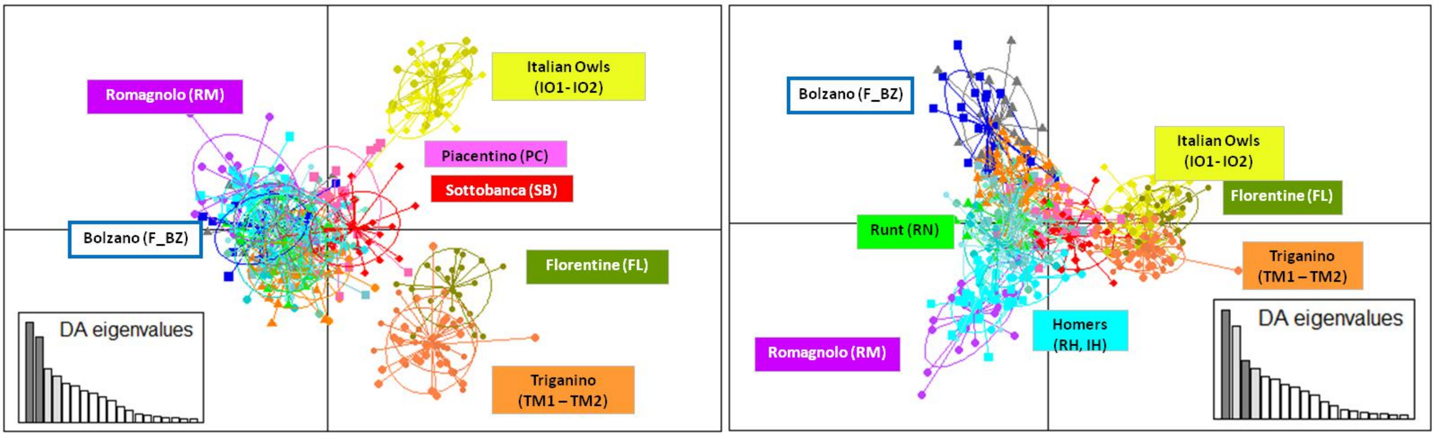

Romagnolo (RM)
Italian Owls (IO1- IO2)
Piacentino (PC)
Sottobanca (SB)
Bolzano (F_BZ)
Florentine (FL)
DA eigenvalues
Triganino (TM1 – TM2)
Bolzano (F_BZ)
Italian Owls (IO1- IO2)
Florentine (FL)
Runt (RN)
Triganino (TM1 – TM2)
Homers (RH, IH)
Romagnolo (RM)
DA eigenvalues


**(b)**

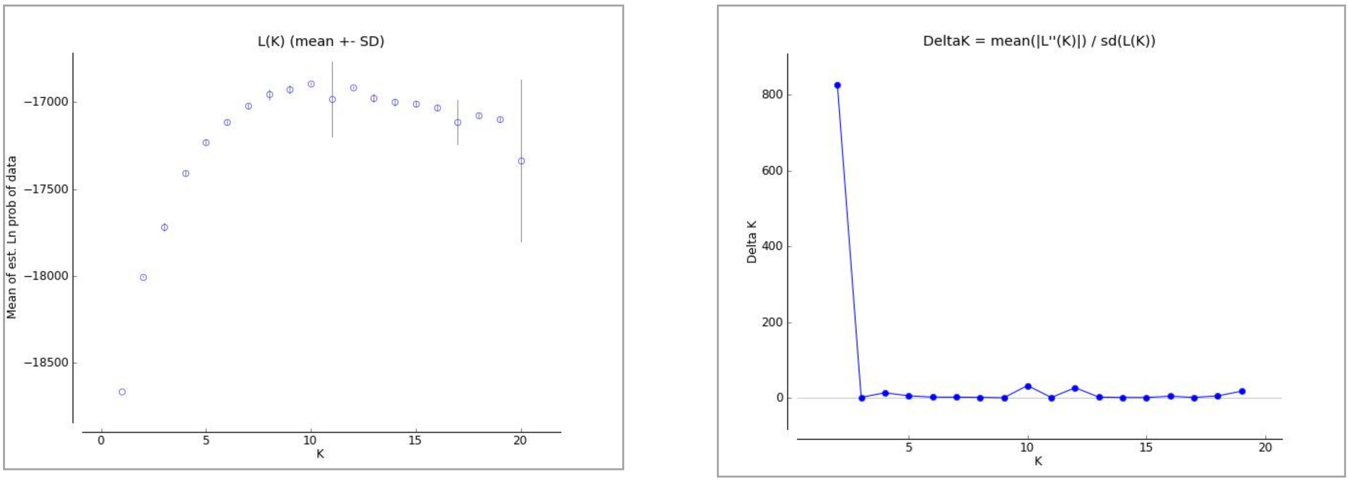

L(K) (mean +- SD)
Mean of est. Ln prob of data
K
DeltaK = mean(|L''(K)|) / sd(L(K))
Delta K
K


**(c)**

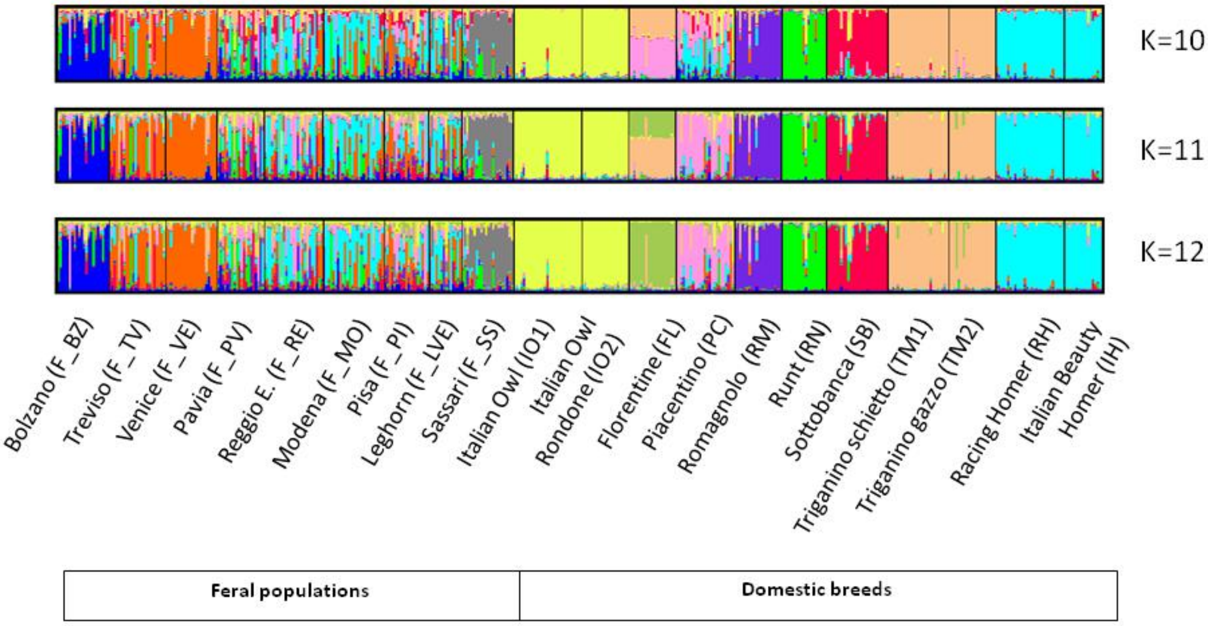

K=10
K=11
K=12
Bolzano (F_BZ)
Treviso (F_TV)
Venice (F_VE)
Pavia (F_PV)
Reggio E. (F_RE)
Modena (F_MO)
Pisa (F_PI)
Leghorn (F_LVE)
Sassari (F_SS)
Italian Owl (IO1)
Italian Owl Rondone (IO2)
Florentine (FL)
Piacentino (PC)
Romagnolo (RM)
Runt (RN)
Sottobanca (SB)
Triganino schietto (TM1)
Triganino gazzo (TM2)
Racing Homer (RH)
Italian Beauty Homer (IH)
Feral populations
Domestic breeds

**ESM 1** Discriminant Analysis of Principal Components (DPCA) and Bayesian computations calculated on feral and on the Italian domestic lineages considered in Bigi et al. (2016). a) In the DAPC plot, the breeds that were supposed not to be involved in the constitution of feral populations (IO1, IO2, FL, TM1, TM2) cluster separately from ferals and from the remaining domestic breeds. b) The best clustering was chosen through ΔK and mean likelihelihood computation at K=10 as the peak at K=12 has been probably produced by a high standard error value in the likelihood computation. c) The bar plot (K = 12) obtained in STRUCTURE shows a clear differentiation between feral and domestic lineages although some domestic components can be found in urban individuals from Lombardy, Emilia-Romagna and Tuscany. The plot at K = 8 and K = 10 were chosen to describe any significant difference in the previous splits.

**ESM 2** F*st* values computed among feral populations. Bold values are not significant after Bonferroni correction ($p > 0.05$).

| | F_TV | F_VE | F_PV | F_RE | F_MO | F_PI | F_LE | F_SS |
|---|---|---|---|---|---|---|---|---|
| F_BZ | 0.06449 | 0.09292 | 0.06101 | 0.05229 | 0.06462 | 0.06049 | 0.06034 | 0.07565 |
| F_TV | | 0.02392 | 0.01926 | 0.02438 | 0.03182 | 0.01591 | 0.03551 | 0.05027 |
| F_VE | | | 0.07514 | 0.06257 | 0.07002 | 0.0667 | 0.06177 | 0.08131 |
| F_PV | | | | **0.01444** | 0.02400 | **0.00618** | 0.02484 | 0.04671 |
| F_RE | | | | | **0.00376** | **0.00996** | **0.00331** | 0.04422 |
| F_MO | | | | | | **0.01359** | **0.00402** | 0.05106 |
| F_PI | | | | | | | 0.02076 | 0.04383 |
| F_LE | | | | | | | | 0.05029 |

(a)

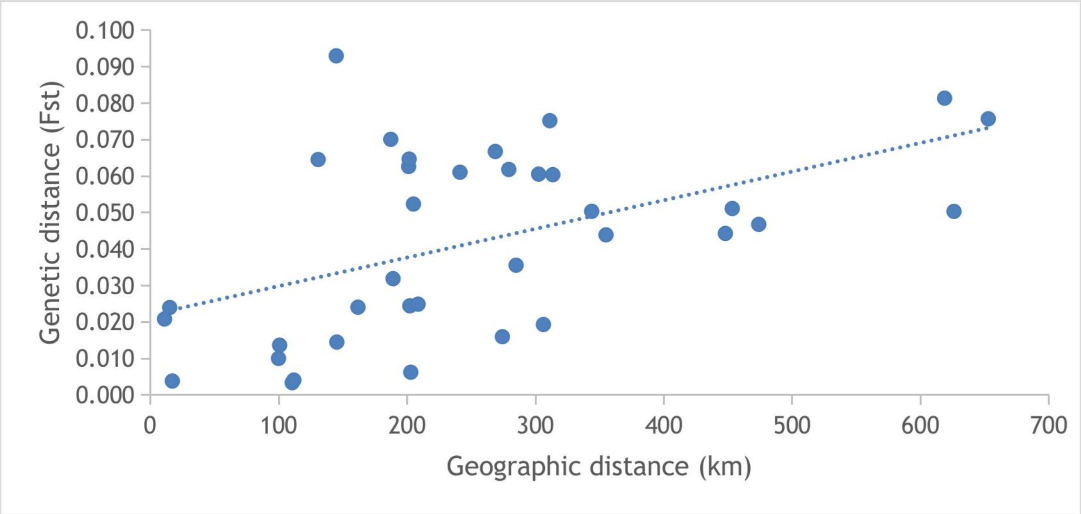


(b)

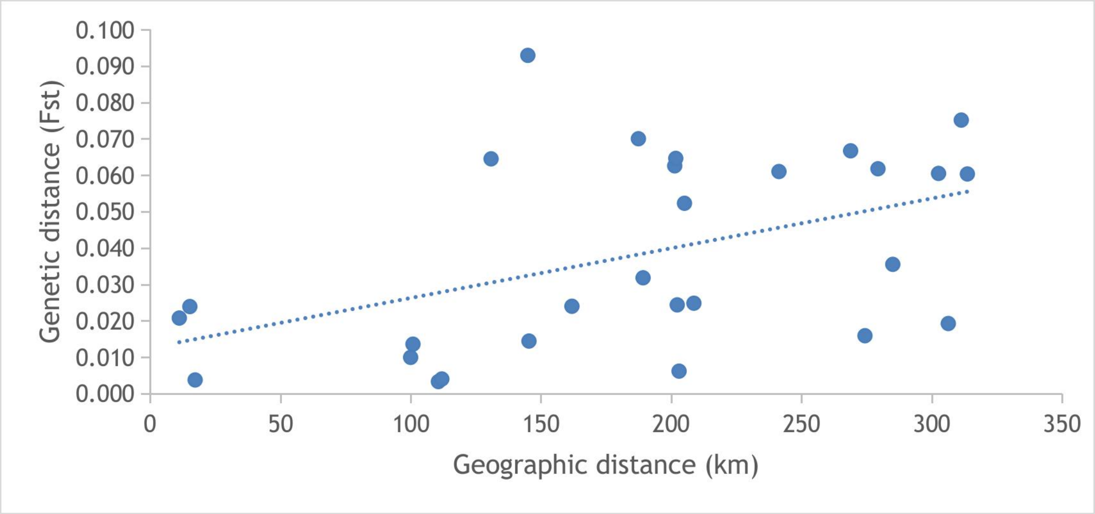


**ESM 3** Relationship between geographic and genetic distance in all feral populations (a) and only in peninsular populations (b). A significant correlation was found in both the analyses (a: $r = 0.50$, $P = 0.03$; b: $r = 0.46$, $P = 0.01$; Mantel test). Dotted lines: fitted lines of the linear model *genetic distance ~ geographic distance.*

**(a)**

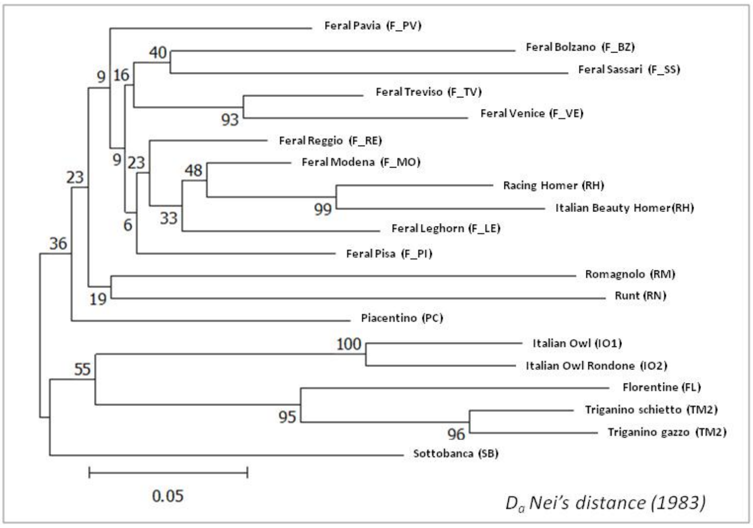

Feral Pavia (F_PV)
Feral Bolzano (F_BZ)
Feral Sassari (F_SS)
Feral Treviso (F_TV)
Feral Venice (F_VE)
Feral Reggio (F_RE)
Feral Modena (F_MO)
Racing Homer (RH)
Italian Beauty Homer(RH)
Feral Leghorn (F_LE)
Feral Pisa (F_PI)
Romagnolo (RM)
Runt (RN)
Piacentino (PC)
Italian Owl (IO1)
Italian Owl Rondone (IO2)
Florentine (FL)
Triganino schietto (TM2)
Triganino gazzo (TM2)
Sottobanca (SB)
0.05
$D_a$ Nei's distance (1983)


**(b)**

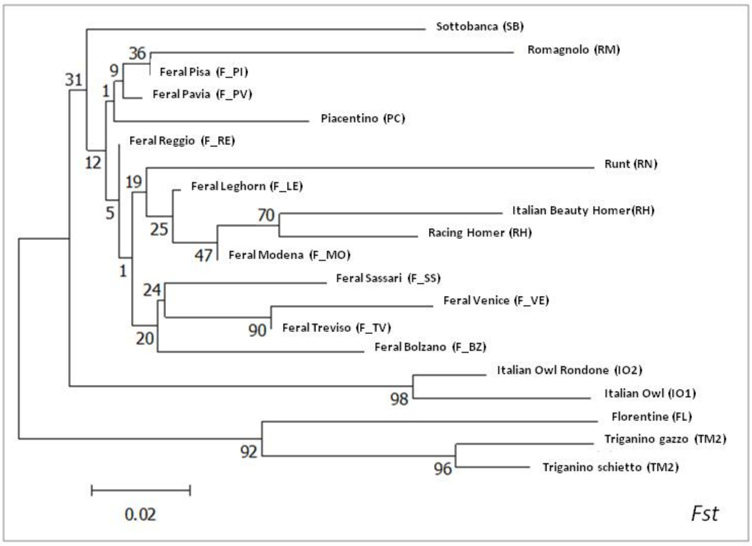

Sottobanca (SB)
Romagnolo (RM)
Feral Pisa (F_PI)
Feral Pavia (F_PV)
Piacentino (PC)
Feral Reggio (F_RE)
Runt (RN)
Feral Leghorn (F_LE)
Italian Beauty Homer(RH)
Racing Homer (RH)
Feral Modena (F_MO)
Feral Sassari (F_SS)
Feral Venice (F_VE)
Feral Treviso (F_TV)
Feral Bolzano (F_BZ)
Italian Owl Rondone (IO2)
Italian Owl (IO1)
Florentine (FL)
Triganino gazzo (TM2)
Triganino schietto (TM2)
0.02
Fst


**(b)**

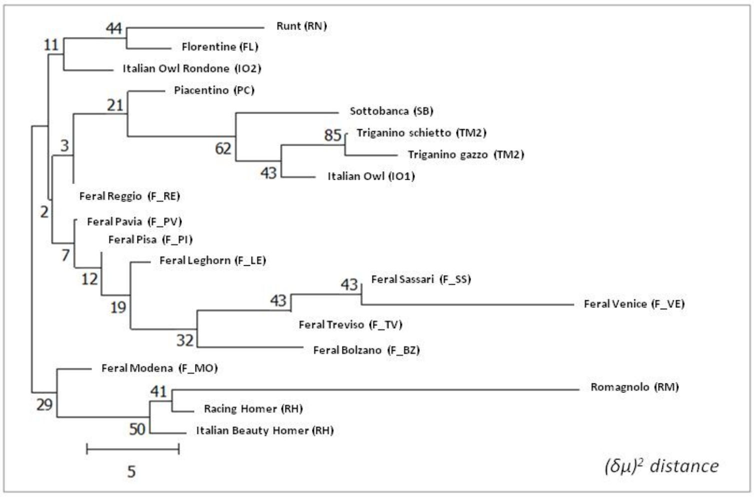

Runt (RN)
Florentine (FL)
Italian Owl Rondone (IO2)
Piacentino (PC)
Sottobanca (SB)
Triganino schietto (TM2)
Triganino gazzo (TM2)
Italian Owl (IO1)
Feral Reggio (F_RE)
Feral Pavia (F_PV)
Feral Pisa (F_PI)
Feral Leghorn (F_LE)
Feral Sassari (F_SS)
Feral Venice (F_VE)
Feral Treviso (F_TV)
Feral Bolzano (F_BZ)
Feral Modena (F_MO)
Romagnolo (RM)
Racing Homer (RH)
Italian Beauty Homer (RH)
5
$(\delta\mu)^2$ distance

**ESM 4** Phylogenetic trees of domestic and feral lineages drawn using three different distance models. a) $D_a$ distance (Nei 1973); b) *Fst;* c) $(\delta\mu)^2$ distance. Only nodes characterized by bootstrap values higher than 50 can be considered significantly supported.
